# Biochemical Reduction of the Topology of the Diverse WDR76 Protein Interactome

**DOI:** 10.1101/650028

**Authors:** Gerald Dayebgadoh, Mihaela E. Sardiu, Laurence Florens, Michael P. Washburn

## Abstract

A hub protein in protein interaction networks will typically have a large number of diverse interactions. Determining the core interactions and the function of such a hub protein remains a significant challenge in the study of networks. Proteins with WD40 repeats represent a large class of proteins that can be hub proteins. WDR76 is a poorly characterized WD40 repeat protein with possible involvement in DNA damage repair, cell cycle progression, apoptosis, gene expression regulation, and protein quality control. WDR76 has a large and diverse interaction network that has made its study challenging. Here, we rigorously carry out a series of affinity-purification coupled to mass spectrometry (AP-MS) to map out the WDR76 interactome through different biochemical conditions. We apply AP-MS analysis coupled to size exclusion chromatography to resolve WDR76-based protein complexes. Furthermore, we also show that WDR76 interacts with the CCT complex via its WD40 repeat domain and with DNA-PK-KU, PARP1, GAN, SIRT1, and histones outside of the WD40 domain. An evaluation of the stability of WDR76 interactions led to focused and streamlined reciprocal analyses that validate the interactions with GAN and SIRT1. Overall, the approaches used to study WDR76 would be valuable to study other proteins containing WD40 repeat domains, which are conserved in a large number of proteins in many organisms

## Introduction

Hub proteins within protein interaction networks are characterized by their participation in large numbers of interactions. Hub proteins have therefore been of great interest as dissection and characterization of their multiple interactions remains challenging (Patil et al, 2010; Tsai et al, 2009; Uchikoga et al, 2016). One such family of hub proteins include proteins containing the WD40 repeat domain, which is the most interacting domain in *S. cerevisiae* and the fourth most abundant domain in the human proteome (Stirnimann et al, 2010). The WD40 repeat domain is typically characterized by the presence of 4-8 repeats of 40-60 amino acids and ending with a tryptophan-glutamic acid (W-D) dipeptide (Higa et al, 2006; Stirnimann et al, 2010). WD40 repeat proteins generally carry at least one such domain, whose structure has been solved and characterized as a tridimensional β propeller. Due to the promiscuity of the WD40 repeat domain, the biological functions of members of this family are diverse with potential applications in therapeutics (Schapira et al, 2017; Zou et al, 2016).

WD Repeat-Containing Protein 76 (WDR76) is a predicted member of the WD40-repeat-containing domain superfamily and has been involved in a variety of distinct biological processes. First described in *S. cerevisiae*, the WDR76 homologue, named YDL156w/CMR1, has been shown to affinity purify with histones (Gilmore et al, 2012), bind UV-damaged DNA (Choi et al, 2012), co-express with genes involved in DNA metabolism (Abu-Jamous et al, 2013), be involved in DNA replication stress (Gallina et al, 2015), and be recruited to coding regions and promote transcription (Jones et al, 2016). The mouse homologue strongly binds H3K27ac and H3K4me3 in mouse embryonic stem cells (Ji et al, 2015). Studies aimed at unveiling novel DNA methylation readers have shown strong WDR76 binding to 5-(Hydroxy)-methylcytosine (relative to other methylcytosine modifications), hence suggesting involvement of WDR76 in epigenetic transcriptional regulation (Spruijt et al, 2013). In a proteomic screen for proteins associated with human mitotic spindles, Sauer *et al.* suggested that WDR76 was a spindle-binding protein (Sauer et al, 2005). However, a later study instead found WDR76-association with chromosomes but not mitotic spindles and showed that depletion of WDR76 resulted in slight mitotic delay and broad metaphase plate (Rojas et al, 2012). Despite the possible implication of WDR76 in multiple distinct biological processes, the exact molecular mechanism of how it performs such role remains to be fully elucidated in most of these processes.

Some clarity regarding the potential function for WDR76 originates in an earlier study suggesting that WDR76 is a CUL4-DDB1 ubiquitin ligase associated factor (DCAF) with possible involvement in histone methylation (Higa et al, 2006). DCAFs can be critical substrate receptors in providing specificity to the CUL4-DDB1 ubiquitin ligase system to regulate several important biological processes (Lee & Zhou, 2007). WDR76 has been linked to the CUL4-DDB1 ubiquitin ligase complex in mammals where WDR76 is involved in regulating circadian rhythms (Tamayo et al, 2015). In addition, recently Jeong *et al.* have shown that the CUL4-DDB1-WDR76 ubiquitin ligase complex is vital for the regulation of RAS levels via a polyubiquitination-dependent degradation mechanism required for inhibition of proliferation, transformation, and invasion in liver cancer cells (Jeong et al, 2019). Collectively, these studies support the idea that WDR76 may be an ubiquitin ligase E3 linker protein.

Given the diverse array of biological processes that WDR76 has been linked to in multiple organisms and the possibility that it could play vital roles in disease etiology, maintenance, and progression, determining the protein interaction network of WDR76 is of high importance. To date, this has been done with affinity purification and mass spectrometry (AP-MS) under one experimental condition (Gallina et al, 2015; Gilmore et al, 2016; Spruijt et al, 2013). The largest study analyzing human WDR76 interacting proteins found a diverse array of more than 100 interactions including histones, heterochromatin related proteins, and DNA damage proteins (Gilmore et al, 2016), demonstrating that WDR76 can be considered a hub protein. Of high importance is discerning the strongest/core interactions from the weaker/transient interactions within the WDR76 interaction network. To this end, we pursued a multi-faceted, high-depth, and system-wide AP-MS strategy to capture the global WDR76 interactome under different conditions. Firstly, we mapped the global WDR76 interactome in HEK293T cells by implementing a high salt lysis procedure. Next, we performed size exclusion chromatography to establish the co-fractionation profiles of members of WDR76 interactome. In addition, we used truncation mutants to map interactions to the two dominant domains of WDR76: the non-WD40 domain and the WD40 domain. Our analysis suggested that the WD40 domain of WDR76 interacts mainly with CCT (Chaperonin Containing TCP1 or TriC-TCP-1 Ring Complex), while the non-WD40 domain interacts with DNA-PK-KU, PARP1, SIRT1, and histones. This finding suggests that the N-terminus of WDR76 may be important to confer WDR76-specific functions in human cells. To assess the strength of interactions within the WDR76 interactome, we further performed affinity purifications using three different wash conditions of increasing salt concentration. Topological data analysis of the resulting mass spectrometry dataset showed that the core/strongest members of the WDR76 interactome include lamins, the CCT complex, GAN, and SIRT1. Taken together, we present a detailed workflow for the refinement of the interactome of highly promiscuous proteins like the WD40 repeat proteins.

## Results

### WDR76 Conservation and Links to Human Disease

WDR76 (UniProtKB: Q9H967) is a novel poorly characterized protein member of the WD40 repeat protein family. To understand the evolutionary relatedness between WDR76 homologues in eukaryotes, we carried out phylogenetic analysis on 17 unique sequences of WDR76 homologues downloaded from NCBI (NCBI HomoloGene:38573). MUSCLE alignment (Chojnacki et al, 2017) showed high bootstrap values between WDR76 and its homologues in eukaryotes (ranging between 0.8-1.0). As expected, human WDR76 was closer to its homologues in vertebrates (being closest to WDR76 in the *Macaca mulatta*) than those in non-vertebrates (Fig. 1A). In all 17 homologues of WDR76, the presence of a C-terminal WD40 domain is predicted (Fig. 1B, Supplementary Figure S1). Since no crystal structure of human WDR76 exists, we carried out three-dimensional structure prediction of the WD40 domain using Phyre2, a web-based server (Kelley et al, 2015). In agreement with predictions, the 3-D structure revealed the presence of a coiled β-propeller architecture (estimated at >90% accuracy) containing seven WD40 blades (Fig. 1C).

**Figure 1.**
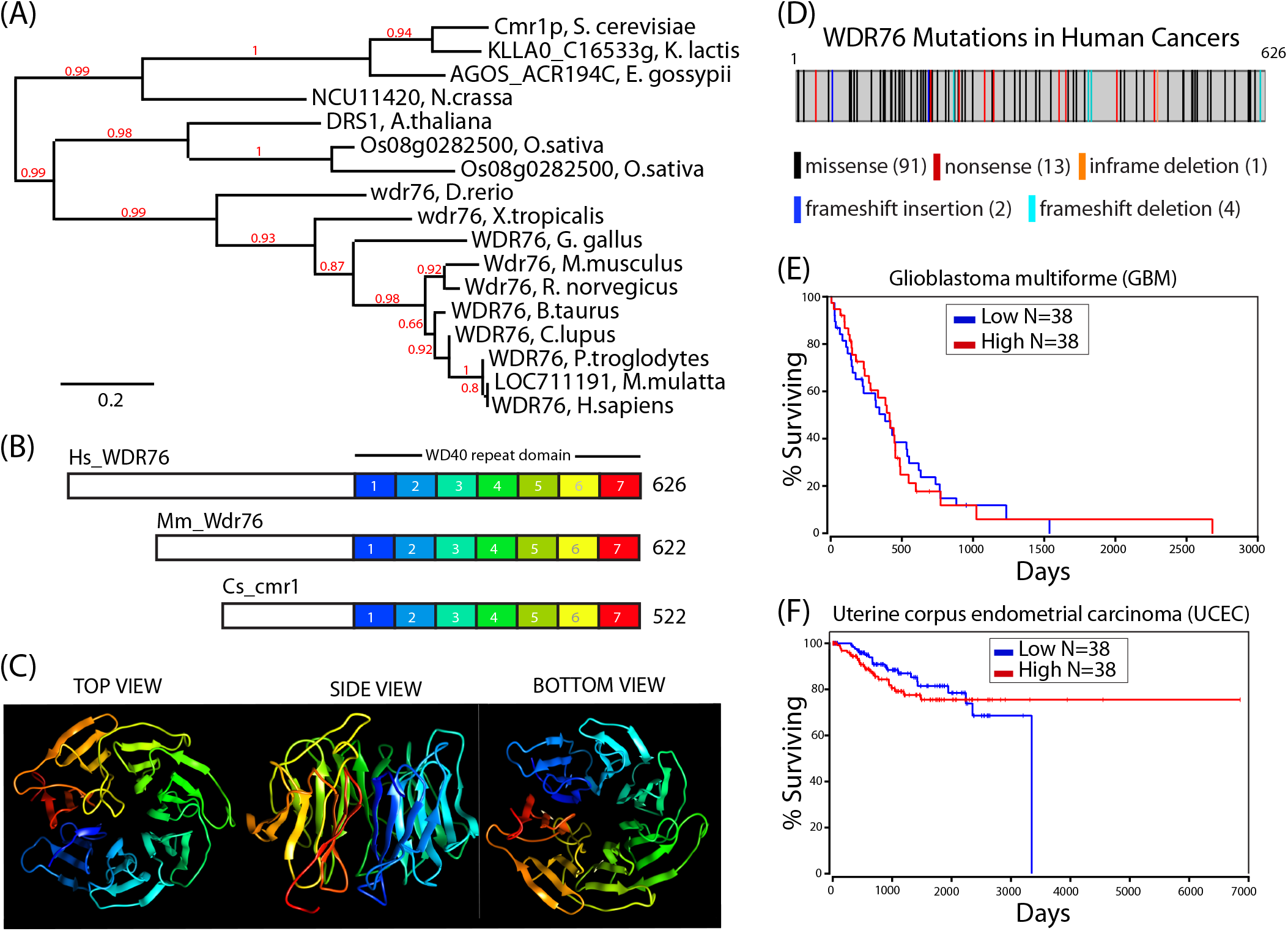
Conservation of WDR76 primary and tertiary structures and disease links. (A) Likelihood Phylogenetic tree among eukaryotes of WDR76 protein homologs. Seventeen WDR76 genes homologs were identified as putative homologues of one another during the construction of HomoloGene. The phylogenetic tree was constructed using www.phylogeny.fr using MUSCLE for alignment, Gblocks for curation, PhyML for Phylogeny and TreeDyn for Tree Rendering (Dereeper et al, 2008). Bootstrap value are shown at nodes (in red). (B) Cartoon of WDR76 homologues in Homo sapiens (Hs_WDR76), Mus musculus (Mm_Wdr76), and Saccharomyces cerevisiae (Cs_cmr1) showing a poorly characterized N-terminal domain and a highly conserved C terminal WD40 domain comprised of 7 WD40 repeats. (C) Prediction of a seven-bladed propeller-like structure for the WD40 domain of WDR76 was generated using Phyre2 (Kelley et al, 2015). The WD40 repeats (blades) are color coded as in (B), withWD1 being the first N-terminal WD40 repeat and WD7 the last C-terminal. (D) Map of 114 WDR76 aberrations reported in COSMIC, including mutations (missense and nonsense), frameshift deletions, frameshift insertions, and an in-frame deletions (Tate et al, 2019). (E-F) Prognostic significance of WDR76 in glioblastoma multiforme and uterine corpus endometrial carcinoma showing correlation of aberrant WDR76 expression with shorter survival.

WDR76 has also been linked to human diseases like Alzheimer’s and cancer. For example, a large whole exome sequencing study searching for rare and ultra-rare variants in Alzheimer’s diseases suggested that WDR76 variants may be linked to the disease (Raghavan et al, 2018). Next, to identify WDR76 aberrations in human cancers, we searched for WDR76-associated mutations in the COSMIC database (Tate et al, 2019) where 110 WDR76 mutations in human cancers were found (Fig. 1D). To further assess the deleterious effects of aberrant WDR76 expression and possible effects on cancer survival in humans, we conducted an OncoLnc database search (http://www.oncolnc.org/) for WDR76. Our search revealed that aberrant expression of WDR76 correlated with shorter survival in some cancers. For example, overexpression and repression of WDR76 in glioblastoma multiforme (GBM) correlated with decreased survival rates (Fig. 1E). Similarly, repression of WDR76 in uterine endometrial carcinoma (UCEC) correlated with shorter survival (Fig. 1F). In addition, the prognostic summary in the Pathology Atlas for WDR76 in The Human Protein Atlas (Uhlen et al, 2017) lists WDR76 as an unfavorable prognostic marker in renal cancer, lung cancer, pancreatic cancer, and liver cancer. These links to cancer are supported by recent evidence of the role of WDR76 in regulating the RAS oncogene (Jeong et al, 2019).

### The Diverse WDR76 Protein Interaction Network

To build a comprehensive interaction network of the human WDR76 interactome, we used the HaloTag affinity purification system (Daniels et al, 2012) coupled with multidimensional protein identification technology (MudPIT) and label free quantitative proteomics analysis (Zhang et al, 2010). Previous affinity purification-mass spectrometry (AP-MS) analyses suggested that both N- and C termini of WDR76 were accessible for tagging and AP-MS analysis (Gallina et al, 2015; Gilmore et al, 2016; Spruijt et al, 2013). However, the placement of an affinity tag can strongly influence the assembly of functional protein complexes (Banks et al, 2018a). Therefore, we first tested which termini of WDR76 could provide better depth into the WDR76 interactome. We transiently expressed N-terminally and C-terminally Halo-tagged WDR76 (Halo-WDR76 and WDR76-Halo, respectively) in HEK293T cells (Fig. S2A). Western blot analysis showed full length Halo-WDR76 and WDR76-Halo were expressed (Fig.S2B; lanes 3 and 4, respectively). In agreement with previous studies from our group (Gilmore et al, 2016) and the Human Protein Atlas (Thul et al, 2017) the N-terminally tagged WDR76 localizes entirely in the nucleus (Fig. S2C). However, the C-terminally tagged WDR76 (WDR76-Halo) showed a more diffuse localization (Fig. S2D). AP-MS analysis revealed that Halo-WDR76 provides a better coverage to the WDR76 interactome (Supplementary Table S1A-D). Thus, HEK293FRT with stable expression of Halo-WDR76 was chosen for subsequent AP-MS mapping of high confidence WDR76 interactions.

We took a comprehensive approach to defining the WDR76 interaction network by purifying Halo-WDR76 from HEK293FRT cells under different biochemical conditions. First, we prepared chromatin-enriched nuclear lysates from HEK293FRT stably expressing of N-terminally tagged Halo-WDR76. Preparation of chromatin-enriched nuclear extracts were conducted in two steps. Nuclear extracts were prepared from cells followed by solubilization of the insoluble pellets in Micrococcal nuclease (Mnase) buffer. Nuclear lysates and chromatin lysates were pooled together to constitute chromatin-enriched nuclear lysates (Fig. 2A). Next, whole cell extracts were prepared by lysis of Halo-WDR76 stable HEK293FRT cells in lysis buffer (Fig.2B). Four replicates of affinity purified Halo-WDR76 versus 4 mock purifications were performed for analysis by label free quantitative proteomics in each approach (Supplementary Table S2). To provide a comprehensive map of the WDR76 interactome, we included our AP-MS data published previously as shown in Fig. 2C) (Supplementary Table S2A and B). Enrichment analysis was performed using QSPEC analysis (Choi et al, 2008) by comparing our purifications relative to mock purifications. Taken together, 115, 89 and 42 WDR76 interactions were significantly enriched in the nuclear extract (Gilmore et al, 2016), chromatin-enriched nuclear extract, and whole cell extract, respectively (Fig. 2A and Supplementary Fig. S3A-C).

**Figure 2.**
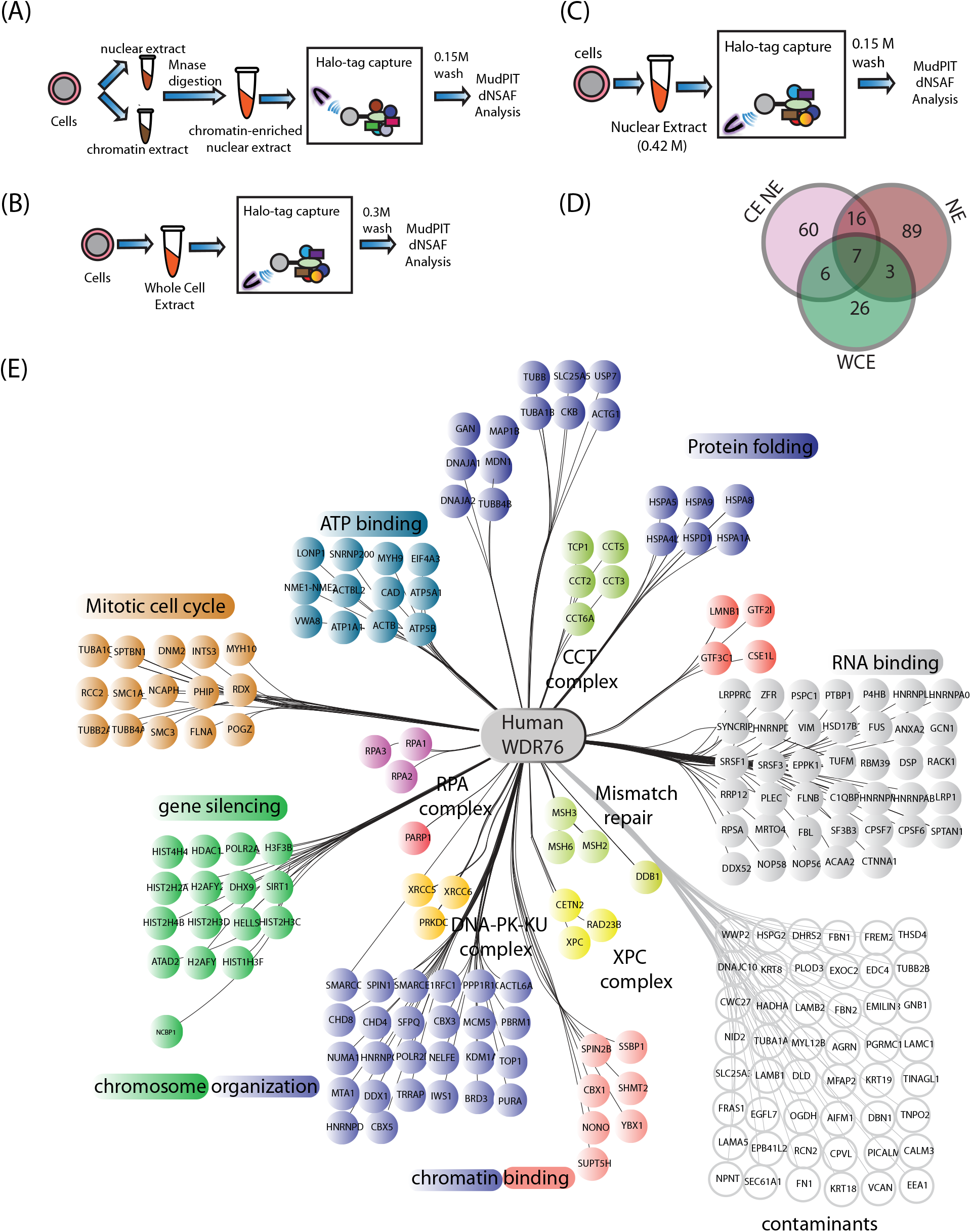
Assembly of WDR76 Protein Interaction Network. Proteins significantly enriched from affinity purifications and quantitative proteomic analysis of Halo-WDR76 from stable HEK293FRT cell lines compared to mock purifications. Four replicates of purifications approach were done (4 controls per experiment) and following QSPEC analysis, cut-off criteria were set at Z≥2, log2FC ≥ 2 or FDR ≤ 0.05. (A-C) Workflows for the AP-MS analysis of chromatin-enriched nuclear extract whole cell extract and nuclear extract for WDR76 interatome mapping. (D) Venn diagram of the significantly enriched proteins from the chromatin enriched nuclear extract (CE NE) dataset, the whole cell extract (WCE) dataset, and the previously published nuclear extract (NE) dataset (Gilmore et al, 2016). (E) Merge of datasets obtained from A-D above.

Next, these three datasets were merged to assemble a global network of human WDR76 interactions (Fig. 2D-E). Functional annotation (gene ontology analysis) of WDR76-assocaited proteins was conducted using EnrichR (Kuleshov et al, 2016). Related proteins were then organized into a WDR76 centralized network that contains proteins involved in protein folding, RNA binding, mitotic cell cycle, gene silencing, chromosome organization, chromatin binding, the XPC complex, the RPA complex, the DNAPK complex, the mismatch repair complex, and the CCT complex (Fig. 2E). This is a broad and diverse spectrum of potential interacting proteins that further supports the role of WDR76 as a hub protein. A total of 207 proteins are represented in Figure 2E. This includes proteins involved in chromatin binding and chromosome organization, such as SPIN2B and CBX1, KDM1A, CBX5, SPIN1, MTA1 and NUMA1, and proteins involved in gene silencing like SIRT1, HELLS and HDAC1. We have previously shown recruitment of WDR76 to sites of DNA damage (Gilmore et al, 2016). Cmr1 also shows preferential binding to uv-damaged DNA (Choi et al, 2012), suggesting that that WDR76 could play a conserved role in the DNA damage recognition or repair process. Many proteins involved in DNA damage repair like XRCC5, XRCC6, DNA-PK, XPC, CETN2, RAD23, RPA1, RPA2 RPA3, MSH2, MSH3, MSH6, DDB1, and PARP1 were also identified (Fig. 2E). In addition, WDR76-deficiency has been linked to defects in the mitotic cell cycle (Rojas et al, 2012), and our data showed significant enrichment of mitotic cell cycle proteins such as RCC2, NCAPH and SMC1A. Next, WDR76 interaction with the chaperonin containing TCP1 or TriC-TCP-1 Ring Complex which has been predicted as an important player in WDR76-mediated protein quality control (Gallina et al, 2015; Gilmore et al, 2016). In addition to the CCT complex, we found significant enrichment the BTB/kelch family protein, GAN which has been linked to giant axonal neuropathy (Bomont et al, 2000). Intriguingly, we also detected a significant enrichment of MAP1B, a known GAN-interacting protein and substrate for GAN-dependent polyubiquitination and proteasomal degradation which plays a vital role in microtubule stability (Allen et al, 2005). Our map of the WDR76 interactome is composed of functionally diverse protein categories. Major challenges in deciphering the biological role of WDR76 include determination of the distinct multi-subunit protein complexes and the core components of this interactome. Reciprocal AP-MS analysis is a very useful approach for the validation of bona fide interactions however it would be cost prohibitive and time prohibitive to carry out reciprocal purifications of all these potential protein interactions. Therefore, different approaches are needed to streamline this list to determine what might be the best candidates for reciprocal purifications, for example.

### Resolving Protein Complexes with Size Exclusion Chromatography

To begin to parse out distinct WDR76 associated protein complexes we analyzed affinity purified Halo-WDR76 from nuclear extracts of stably transfected HEK293FRT cells via size exclusion chromatography coupled to label free quantitative proteomics as shown in Figure 3A. Affinity purified WDR76 was loaded onto a Superose 6 column and 26 fractions of 500μl were collected, digested and analyzed by label free quantitative proteomic analysis (Supplementary Table S3A). The approximate molecular weight of WDR76 is 70kDa, however, the size exclusion fractionation profile of WDR76 showed WDR76-specific peptides above 70kDa, suggesting that WDR76 is assembled into larger complexes. To determine the molecular composition of potential WDR76-associated multiprotein complexes across the 26 fractions from size-exclusion chromatography, we used topological scores (TopS) (Sardiu et al, 2019) to compare protein abundances across all fractions (Fig. 3B and Supplementary Table S3B). We previously showed that topological scores efficiently discriminate strong interaction partners from weak interactions when different affinity purifications are compared (Sardiu et al, 2019). We selected 47 proteins that had TopS score greater than 5 in at least one fraction and were previously identified in Fig. 2 (Figure 3B, Supplementary Table S3C). A large number of distinct potential protein complexes were seen in this analysis. For example, based on protein composition and fractionation profiles, WDR76 co-fractionated with heterochromatin 1 protein members (CBX1, CBX3 and CBX5) in fractions 24-25. A similar pattern was observed with SPIN1 forming a low molecular weight complex with WDR76. WDR76 association with the CCT complex forms a large molecular weight complex of approximately 669 kDa in fraction 17 (Fig. 3B). A potential complex with WDR76, GAN, and MAP1B was seen in fractions 19 and 21 (Fig. 3B). WDR76 was also seen associating with the DNA repair proteins XRCC5, XRCC6, and DNA-PK complex in fraction 19 (Fig. 3B). Other possible WDR76-based complexes include SIRT1, HELLS, and BRD3 (Fig. 3B).

**Figure 3.**
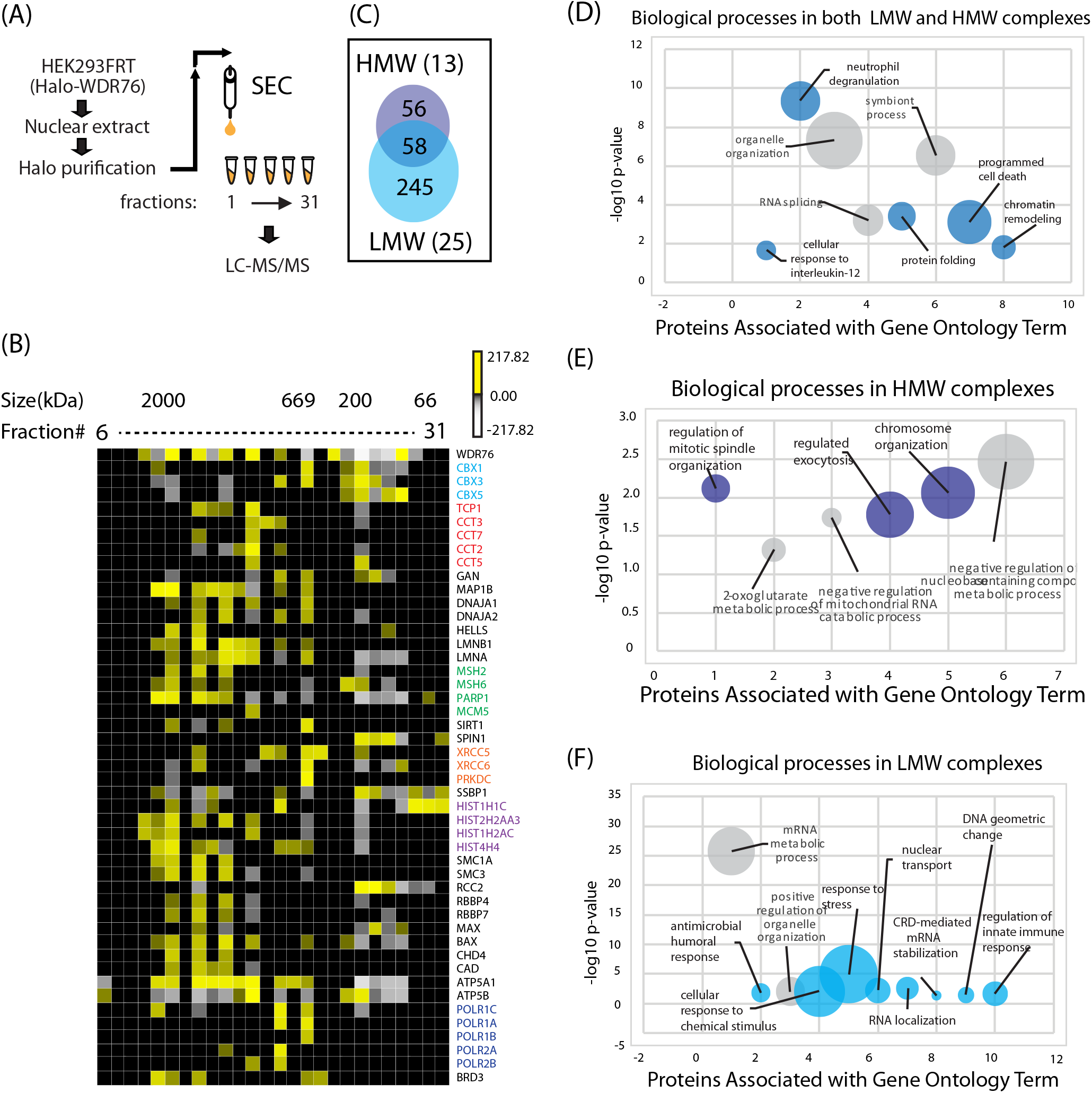
Separation of WDR76 Complexes via Size Exclusion Chromatography. (A) Workflow for preparations of nuclear extract, HaloTag purification, size exclusion chromatography and MudPIT analysis of Halo-WDR76 from HEK293FRT cells stably expressing Halo-WDR76. (B) Topological score (i.e. TopS) was applied to the eluted proteins across 26 fractions. The heat map of 47 protein elution profiles across 26 fractions is represented in here. Only proteins identified as well as in the WDR76 wild-type network with TopS greater than 5 in at least one elution were included in the figure. Yellow color corresponds to high TopS scores whereas grey color corresponds to low TopS values. Black represents the zero values. (C) Venn diagram highlighting proteins with more than 3 peptides in fraction 13 (example of a high molecular weight) and fraction 25 (low molecular weight) from size exclusion chromatography. (D) Gene ontology (GO) terms enriched in high molecular weight fractions. (E) Gene ontology (GO) terms enriched in both high and low molecular weight fractions (F) Gene ontology (GO) terms enriched in low molecular weight fractions.

Based on the corresponding size of each fraction, fractions greater than 200kDa were considered as high molecular weight (HMW) fractions and fractions with less than 200kDa as low molecular weight (LMW) fractions. To gain insight into the biological function of identified WDR76-associated proteins, we selected 2 representative fractions: fraction 13 (HMW) and 25 (LMW) for gene ontology analysis of enriched biological processes. Proteins with more than 2 detected peptides were compared in between fraction 13 and 25 and each fraction contained some unique profile with some overlapping proteins identified in both fractions. Both fractions selected showed enrichment of processes like programmed cell death, protein folding, chromatin remodeling and RNA splicing. The high molecular weight fraction was enriched in processes like regulation of mitotic spindle organization and chromosome organization. In contrast, processes like stress response, regulation of immune response and response to chemical stimuli were enriched in the low molecular weight fraction (Figs. 3D-F, and Supplementary Table S3).

### Domain-specific Interactions of WDR76

Because of high promiscuity of WD40 repeat proteins and the large size of the WD40 repeat protein family (Schapira et al, 2017; Zou et al, 2016), functional specificity of a particular WD40 repeat protein is likely conferred by sequences outside the WD40 domain. Determination of domain-specific interactions of the WDR76 domains could provide clues to the protein interactions important for conferring WDR76-specificity in biological processes. To determine domain-specific interactions of WDR76 we expressed SNAP-tagged full length WDR76 (1-626) and two WDR76 deletion mutants in HEK293T cells. The C-terminal deletion of WDR76 (WDR76Δ) contains residues 1-310, which contains the N-terminus lacking WD40 repeats, and the N-terminal deletion of WDR76 (WDR76Δ’) contains residues 311-626, which contains the domain with the WD40 repeats (Fig. 4A). Full length WDR76 and deletion mutants were affinity purified and analyzed by MudPIT and label free quantitative proteomics.

**Figure 4.**
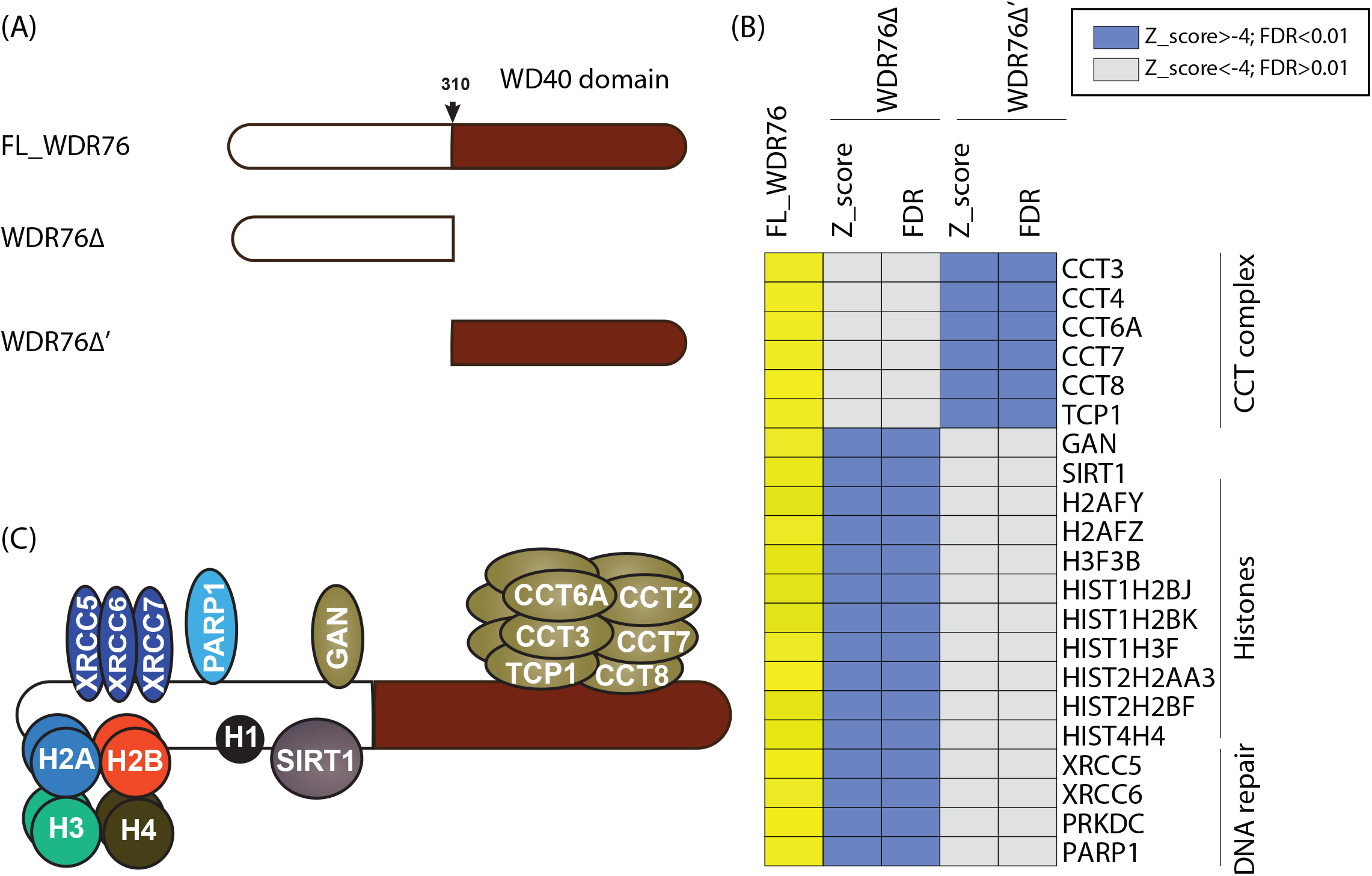
Domain-specific interactions of WDR76. (A) Schematic illustration of full length WDR76 and deletion mutants generated: FL_WDR76, WDR76Δ (residue 1-310) and WDR76Δ’ (residue 311-626). (B) Heat map showing significantly perturbed WDR76 associations resulting from N- and C-terminal domain deletion of WDR76. SNAP-tagged constructs of full length WDR76 and deletion mutants were transiently expressed in HEK293T cells, followed by AP-MS analysis. Q-spec statistics was conducted in 3 replicates of the experiments versus 3 mock purifications. Stringent cutoff values of Z-score≥4 and FDR≤ 0.01 were set to discriminate relevant changes. (C) Model of domain-specific binding of WDR76. We present binding of the C-terminal WD40 domain to the CCT complex and the N-terminal binding of histones, GAN, SIRT1, DNA-Pk-KU complex and PARP1.

In agreement with earlier observations, full length WDR76 showed interaction with the CCT complex, histones, the NAD-dependent deacetylase SIRT1, GAN, DNA-Pk-KU and PARP1 (Fig. 4A and Supplementary Table 4). Interestingly, deletion of the C-terminal WD40 abrogates interaction of WDR76 with the CCT complex but not the interaction with SIRT1, GAN, Histones PRKDC, XRCC5, XRCC6, and PARP1, for example (Fig 74. and Supplementary Table 4). On the other hand, deletion of the N-terminal of WDR76 showed significant interaction with the CCT complex and a significant decrease of WDR76 interaction with SIRT1, GAN, Histones PRKDC, XRCC5, XRCC6, and PARP1 (Fig. 4B, Supplementary Table 4). This finding suggests that the WDR76 interaction of the CCT complex occurs via the WD40 repeat domain of WDR76 while SIRT1, GAN, Histones PRKDC, XRCC5, XRCC6, and PARP1 interact with WDR76 via outside of the WD40 domain (Fig. 4C).

### Evaluation of relative stabilities of proteins the WDR76 interactome

Given the diversity of our initial WDR76 interaction map, we sought to determine the most stable interactions of WDR76 as this could serve as a clue to further information on the functional role of WDR76 in human cells. To biochemically determine the most stable WDR76 interactions, we adopted an AP-MS approach wherein wash steps of the HaloTag® Mammalian Pull-Down protocol were replaced by high salt wash buffer containing increasing NaCl concentrations. We prepared whole cell lysates at near-physiological salt conditions from stable Halo-WDR76 expressing HEK293FRT cells. During affinity purification, however, the wash steps of the affinity purification protocol were replaced by high salt buffer containing 0.5M or 0.75M or 1.0M NaCl (Fig. 5A). WDR76 isolates were analyzed by MuDPIT following the different wash conditions (Fig. 5B-D, Supplementary Table S5A). In addition to unveiling high stability interactions, such salt-resistance evaluations can greatly decrease non-specific background interactions (Joshi et al, 2013).

**Figure 5.**
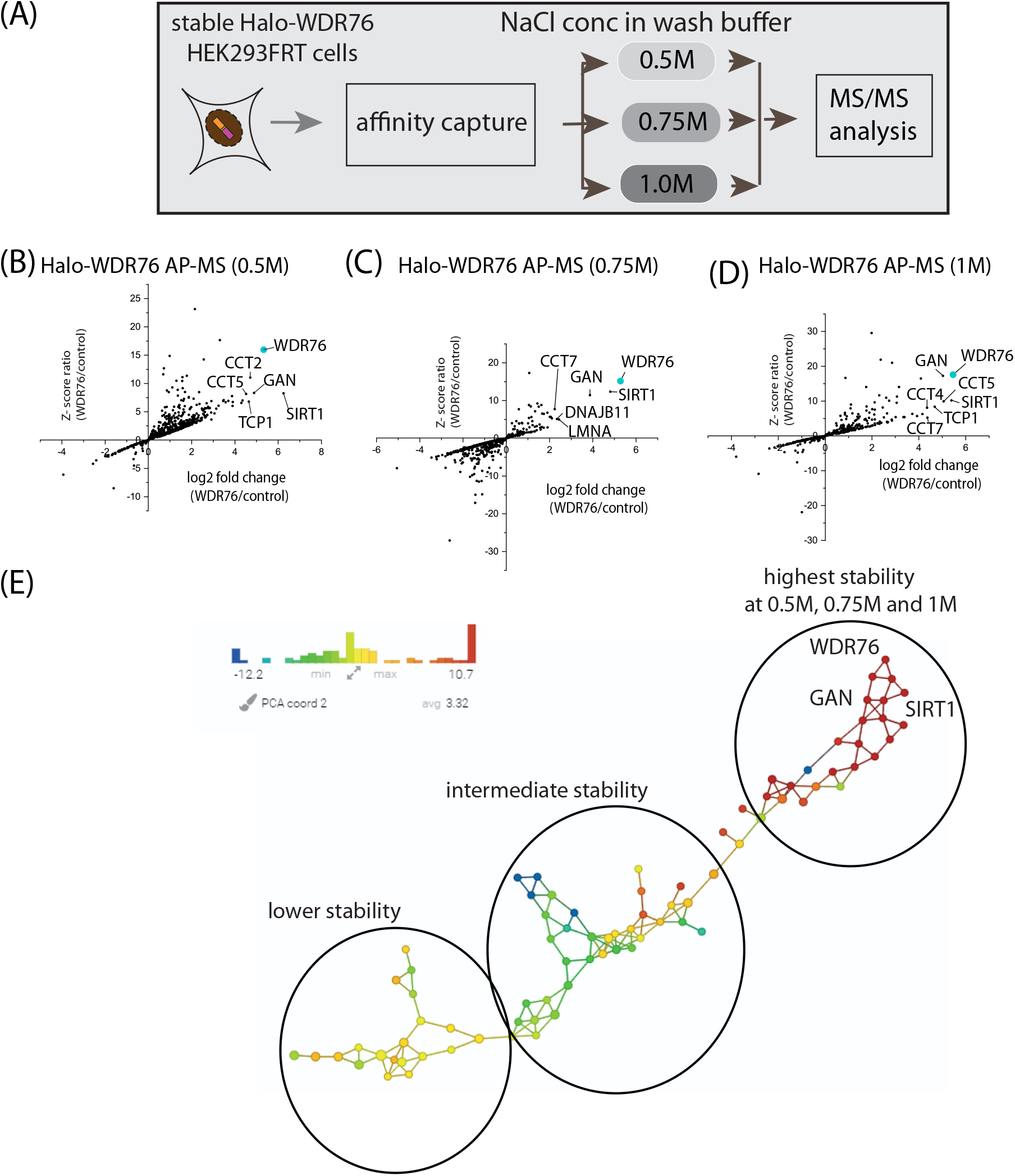
Analysis of Salt Resistant WDR76 Interactions. (A) Workflow showing the preparation of whole cell lysates, Halo-purification and MudPIT of WDR76 isolates from stable Halo-WDR76 HEK293FRT cells. Scatter plot showing AP-MS and significantly enriched interactions in 3 replicates Halo-purifications of WDR76 from stable Halo-WDR76 expressing cells (relative to mock Halo-purifications) at (B) 0.5M (C) 0.75M and (D) 1.0M NaCl wash salt concentrations. (E) TDA was performed on 115 proteins using spectral counts. All the nodes in the network are colored based on metric PCA coordinate 1. Variance normalized Euclidean metric was used with two filter functions: Neighborhood lens 1 and Neighborhood lens2 (resolution 30, gain 3.0x) using the Ayasdi platform. Three main clusters were detected corresponding to the three concentrations used in the analysis. (F) Histogram showing the top 20 WDR76-associated proteins obtained at 1.0M NaCl wash conditions.

To determine the tightest interactions and topology of our protein interaction network, we selected the most enriched (115) WDR76 interactions at 0.5M, 0.75M and 1.0M NaCl wash buffer conditions and conducted topological analysis (Sardiu et al, 2019) (Fig. 5E, Supplementary Table S5). Above 0.5M NaCl wash buffer conditions, the WDR76 interactome composed mainly of lamins, the CCT complex, GAN and SIRT1 and increasing to 1.0 NaCl wash conditions had a minimal impact on these interactions (Fig. 5B-D). The top 17 most stable WDR76 interactions include the CCT complex, the emerin complex (LMNA, LMNB1 and LMNB2), SIRT1, GAN and ECI1 (Fig. 5B-D, Supplementary Figure S4A). Gene ontology (GO) term analysis of enriched biological processes show processes like protein folding and regulation of protein stability (Supplementary Figure S4B). Based on these results, the very diverse WDR76 interactome was further refined to reveal the core components which persist 1.0M wash buffer conditions. As described earlier, it would be time consuming and expensive to carry out reciprocal purifications and label free quantitative proteomic analysis of all 207 potential WDR76 reciprocal associated proteins. Our approach further untangles the WDR76 interactome and ranks proteins into high stability, intermediate stability and low stability members (Fig. 5E). As a hub protein, the salt persistence assay above, have further refined the WDR76 interactome. Summarily, the WDR76 interactome comprises transient or less stable (HELLS, for example) and more stable interacting partners (SIRT1 and GAN for example).

### Expanded WDR76 Centered Protein Interaction Network

To further confirm the proteins identified in our initial AP-MS analyses as bona fide WDR76 interacting proteins, we selected two of the most stable interacting proteins and one weaker interaction for reciprocal validation by AP-MS analysis. We individually and transiently expressed N-terminally HaloTagged GAN, HELLS and SIRT1 HEK293T cells. Following AP-MS analyses, the interactomes of GAN, HELLS and SIRT1 were mapped by MudPIT and label free quantitative proteomic analysis (Supplementary Table S6). In agreement with our earlier data, WDR76 copurified with endogenous GAN, HELLS and SIRT1 (Fig. 6A; Supplementary Table S6A). Reciprocal analysis of three biological replicates of GAN, HELLS and SIRT1 showed copurification of endogenous WDR76 with all three proteins (Fig. 6B, Supplementary Table S6A). In these purifications, WDR76 had the highest enrichment in Halo-GAN, followed by Halo-SIRT1, with Halo-HELLS having the lowest fold change and Z-score for endogenous WDR76 (Fig. 6B). This correlates well with the salt stability results where GAN and SIRT1 are amongst the most stable interactions with Halo-WDR76 (Fig. 5).

**Figure 6.**
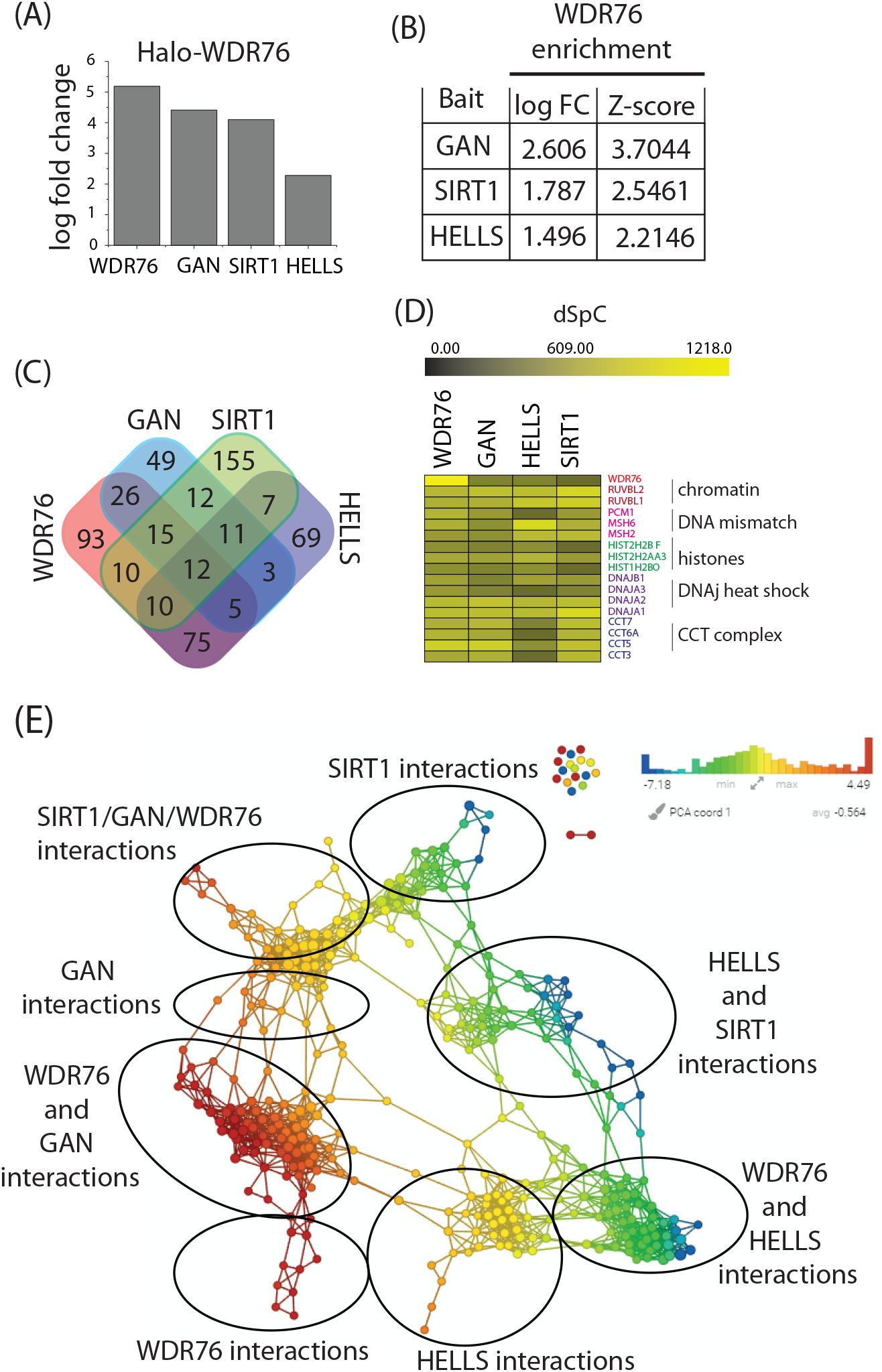
Assembly of Expanded Interactome of WDR76 Interacting Proteins. Reciprocal AP-MS analysis of WDR76 interactions: Case of GAN, HELLS and SIRT1. (A) Histogram depicting co-purification of endogenous GAN, HELLS and SIRT1 with WDR76 (bars show log fold change enrichment compared to control) detected by AP-MS analysis of WDR76. (B) Table showing co-purification enrichment values for endogenous WDR76 (presented as log fold change and Z-score with respect to control) in HaloTag purifications of GAN, HELLS and SIRT1 respectively. Three replicates of HEK293T cells were transiently transfected with each with Halo-GAN, Halo-HELLS or Halo-SIRT1 expressing vectors followed by Halo-purification and AP-MS analysis. (C) Venn diagram comparing significantly enriched interaction obtained WDR76, GAN, HELLS and SIRT1 purifications (Z-score≥2, FDR≤0.05). (D) Heatmap showing of a subset of WDR76 shared with the interactomes of WDR76, GAN, HELLS and SIRT1. (E) Topological network showing distinct interactomes of WDR76, GAN, HELLS and SIRT1 (analysis was done in 3 replicates).

Comparative analysis of the interactomes of WDR76, GAN, and HELLS and SIRT1 showed that each protein had distinct interaction network which only overlap slightly (Fig. 6C). All baits used showed interaction with the CCT complex (CCT3, CCT5, CCT6A and CCT7), DNAJ heat shock proteins (DNAJA1, DNAJA2, DNAJA3 and DNAJB1), DNA mismatch proteins (MS2 and MSH6), chromatin-associated proteins and histones H2A and H2B (Fig. 6D). We next built a TDA network using significantly enriched protein list in each bait. This network separated into eight major areas of WDR76 interactions, WDR76 and GAN interactions, GAN interactions, SIRT1/GAN.WDR76 interactions, SIRT1 interactions, HELLS and SIRT1 interactions, WDR76 and HELLS interactions, and HELLS interactions when moving clockwise from WDR76 interactions (Fig. 6E). The arrangement of this network suggests that WDR76 and GAN share a significant network that is distinct from a SIRT1, GAN, and WD76 network, for example (Fig. 9E). In addition, WDR76 and HELLS share a distinct portion of the network as does HELLS and SIRT1 (Fig. 6E). The binary nature of the WDR76 interaction with each of the 3 preys chosen for validation (GAN, HELLS and SIRT1) confirms WDR76 as a putative hub whose interaction with each prey is important for a different biological function.

## Discussion

WDR76 and its homologues have not only maintained the β-propeller WD40 repeat domain structure but based on experiments from different groups biological functions have been conserved as well including DNA damage recognition and histone binding (Gilmore et al, 2016; Tkach et al, 2012). Cmr1 has been shown consistently shown to bind DNA damage and DNA-metabolic processes with significant H2A serine 129 phosphorylation enrichment in MMS treated cmr1Δ cells (Tkach et al, 2012). Also, human WDR76 is recruited to sites of DNA damage in living HEK293T cells (Gilmore et al, 2016). These findings suggest that WDR76 is a DNA damage response protein. Although recruitment of WDR76 has not been studied in other organisms, observed light-and uv-induced expression of Wdr76 in mouse and zebrafish confirms the protective role of WDR76 and its homologues DNA damage stress response and genome integrity maintenance (Banks et al, 2018a; Gallina et al, 2015; Gavriouchkina et al, 2010; Tamayo et al, 2015). Secondly, both cmr1 and WDR76 show histone binding in vivo (Gilmore et al, 2016; Gilmore et al, 2012) and proteomic analyses of cmr1 interaction profile showed enrichment of histone-related processes which were significantly depleted in histone H4 deficient cells (Gilmore et al, 2012)}. In human histone H1 depletion results in significant WDR76 deletion in embryonic stem cell (Sancho et al, 2008). Although cmr1 promotes transcription genome-wide (Jones et al, 2016), the role of WDR76 in transcription is not yet clear. Recent reports showing the importance of WDR76 in protein quality control warrant clearer understanding of the role WDR76-based CUL4-DDB1 complexes in these processes (Gallina et al, 2015; Gilmore et al, 2016).

Prior to this study, data on the biochemical composition, dynamics and stability of the human WDR76 interactome was limited (Gallina et al, 2015; Gilmore et al, 2016; Spruijt et al, 2013). Our study used affinity purification coupled mass spectrometry and quantitative label free proteomics to map out a high confidence WDR76 interactome in human cells. Using affinity purification and size exclusion chromatography coupled to mass spectrometry, we determined that WDR76 forms multi-subunit protein complexes. For example, the CCT complex associates with the C-terminal WD40 domain of WDR76. The CCT complex has been previously shown to promote folding of the WD40 repeat domain (Miyata et al, 2014). Surprisingly our studies show that WDR76 interacts with CCT via the WD40 domain suggesting that CCT may be important for folding of the WD40 repeat domain in WDR76. In addition, the amino acid residues outside the WD40 repeats interact with SIRT1, GAN, histones and DNAPk-KU complex. Biological specificity of WD40 proteins is largely determined by sequences outside the WD40 domain (van Nocker & Ludwig, 2003). This suggests that WDR76 interaction with SIRT1, GAN, histones and DNAPk-KU could be important for conferring its biological specificity.

Our salt persistence evaluation further refines the WDR76 interactome thereby revealing the strongest interactors and eliminating weak/transient interacting partners of WDR76. Initial AP-MS analysis on cell lysates prepared at 0.42M NaCl containing lysis buffer conditions, we identified 207 high confidence WDR76-binding proteins. Using high salt wash conditions, we determine that GAN, SIRT1, and CCT (Chaperonin Containing TCP1 or TriC-TCP-1 Ring Complex) are the most stable interactors of WDR76. We also determine the weak WDR76 interactions (HELLS for example) do not persist our 1M NaCl wash conditions. GAN, otherwise known as gigaxonin, is a BTB/kelch family protein linked to giant axonal neuropathy (Bomont et al, 2000) and GAN is also involved in the degradation of MAP1B, a process important for neuronal survival (Allen et al, 2005). SIRT1, sirtuin1, is an NAD-dependent deacetylase plays roles in a large number of areas of biology including glucose homeostatis (Rodgers et al, 2005), cell survival (Yeung et al, 2004), and activators of sirtuins have been linked to extended lifespans in yeast (Howitz et al, 2003). These core interactions with GAN and SIRT1 further suggest that WDR76 plays important roles in critical biological processes that warrants further study.

In this body of work, we further present the challenge of studying a protein like WDR76 that has many interactions that from a diverse series of biological pathways. Hub proteins (Patil et al, 2010; Tsai et al, 2009; Uchikoga et al, 2016), like WDR76, remain a challenge to study given these large number of interactions. Here we devised and implemented methods and approaches to reduce the complexity of the WDR76 protein interaction network to determine the strongest interacting proteins that we could then validate using reciprocal purifications. It would have been far more expensive to carry out reciprocal purifications of all the possible WDR76 associated proteins. By using biochemical approaches, like the use of increasing salt concentrations to test interaction stability, we were able to determine proteins that were more likely to pull down WDR76 in a reciprocal purification. This approach is generally valuable for interactome studies and streamlines the process of determining which prey proteins to evaluate as bait proteins to test interaction reciprocity.

## Material and Methods

### Materials

Magne® HaloTag® beads (Promega, G7281) and SNAP-Capture Magnetic Beads (S9145) were from Promega (Madison, WI) and New Englamd Biolabs, respectively. HaloTag® TMRDirect™ Ligand (Promega, G2991) was from Promega (Madison, WI). AcTEV™ Protease (12575015) and PreScission Protease (27-0843-01) were from Thermo Fisher Scientific and GE Health Life Sciences, respectively. HaloTag™ clones of Flexi® vector, pFN21A constructs of GAN (FHC25786), SIRT1 (FHC23876) and HELLS (FHC05413) were from the Kazusa DNA research institute (Kisarazu, Chiba, Japan). SNAP-FLAG pcDNA5 Sgfl/PmeI plasmid was described earlier (Banks et al, 2018b). Flp-In™-293 (HEK293FRT) cells were from ThermoFisher Scientific (Waltham, MA) and were used for all stable transfection-based studies. HEK293T cells (ATCC® CRL11268™) were from ATCC (Manasses, VA) and were used for all transient transfection-based studies. Rabbit anti-HaloTag® polyclonal antibody (G9281) was from Promega. Mouse anti tubulin monoclonal antibody (66031-1-Ig) was from ProteinTech. IRDye® 680LT labeled goat anti-Mouse (926-68020) and IRDye® 800CW labeled goat anti-Rabbit (926-3211) secondary antibodies were from LI-COR Biosciences.

### Plasmids and Cloning of constructs for transient transfection

Full length WDR76 ORF (purchased used) (Gilmore et al, 2016), was digested cut out using AsiSI and pmeI and subcloned into pcDNA5FRTSgfI/PmeI(Banks et al, 2018b) and SNAP-FLAG pcDNA5 SgfI/PmeI (Banks et al, 2018b)to generate Halo-tagged WDR76 and SNAP-tagged WDR76 pcDNA5/FRT expressing constructs. For generation of SNAP-FLAG SNAP-tagged constructs for expression of WDR76 domain deletion mutants: (WDR76Δ and WDR76Δ’), DNA sequences corresponding to the ORF of were amplified by PCR and subcloned into the SNAP-FLAG pcDNA5 vector. To generate C-terminally Halotagged construct of WDR76 (WDR76-Halo) in pCDNA5, PCR was used to amplify the WDR76 DNA sequence ORF and the PCR product was subcloned into pcDNA5FRT C-Halo construct. All PCR primers used in this study are listed in supplementary data.

### Expression of HaloTag Bait proteins in 293 cells

Flp-In™-293 constitutively expressing HaloTag-WDR76 were generated previously (Gilmore et al, 2016). 3 × 10^8^ cells were used for initial WDR76 AP-MS purifications as described elsewhere(Banks et al, 2018b). For transient expressions, 2 × 10^7^ cells were seeded and maintained at 3737°C in 5% CO_2_ for 24 hours after which 10µg of HaloTag constructs: HaloTag-WDR76 (pCDNA5), HaloTag-GAN (pFN21A), HaloTag-SIRT1 (pFN21A), and HaloTag-HELLS (pFN21A), were used to transfect HEK293T cells. For expression of SNAP-tagged WDR76 and truncation mutants, 10µg of plasmid was also used. Lipofetamine 2000 was used as was used as transfection reagent (according to manufacturer’s instructions). Transfected cells were cultured at 37°C in 5% CO_2_ for 48 hours and harvested for downstream applications.

### Cell culture

Cells were Dulbecco’s Modified Eagle’s medium (DMEM) supplemented with 10% fetal bovine serum (FBS) and 2 mM Glutamax at 37°C in 5% CO_2_. The stable Halo-WDR76 Flp-In™-293 cell line was cultured in Dulbecco’s Modified Eagle’s medium (DMEM) supplemented with 10% fetal calf serum (FCS), 1x penicillin and 1x streptomycin. For transient transfections, HEK293T cell cultures were maintained in Dulbecco’s Modified Eagle’s medium (DMEM) supplemented with 10% fetal bovine serum (FBS) and 2 mM Glutamax at 37°C in 5% CO_2_.

### Preparation of chromatin-enriched nuclear extract

Chromatin-enriched nuclear extracts were prepared in two steps. The first step was the preparation of nuclear extracts using Dignam’s method (Dignam, 1990). Initially, cell pellets were resuspended in 5 cell volumes of buffer A (10mM HEPES pH 7.9, 1.5mM MgCl_2_, 10 mM KCl, 0.5mM DTT) supplemented with protease inhibitor and allowed on ice for 10 minutes to swell cells. Swollen cells were centrifuged at 1000rpm at 4°C for 10minutes. The cells were resuspended in 2 cell volumes of buffer A and lysed mechanically with the loose pestle of a dounce homogenizer. Following >90% cells lysis as detected by trypan blue, the cell lysates were centrifuged at 25000rpm for 20minutes. The supernatant was discarded and the packed nuclear pellet volume (PNPV) was determined. The nuclear pellet was responded in 0.11 × PNPV buffer C (20mM HEPES PH7.9, 25% glycerol, 420mM NaCl, 1.5mM MgCl_2_, 0.2mM EDTA, 0.5mM DTT) supplemented with protease inhibitor cocktail and salt active nuclease and homogenized with 2 strokes in Dounce homogenizer (B or loose pestle). Resuspended nuclei were incubated in buffer C at 4°C for 1 hour. The nuclei were centrifuged at 40000rpm for 1 hour to collect the nuclear extract. Using a dounce homogenizer, the resulting pellet was resuspended in Mnase reaction buffer (50 mM HEPES buffer (pH 7.9), 5 mM MgCl2, 5 mM CaCl2 and 25% (v/v) glycerol)) containing MNase ((micrococcal nuclease (M0247S, New England Biolabs)) and supplemented with protease inhibitor. We incubated the mixture at 4°C for 3hours followed by centrifugation at 40,000xg for 1 hour at 4°C to obtain a chromatin extract (supernatant). To obtain a chromatin-enriched nuclear extract, nuclear and chromatin extracts were pooled together for subsequent procedures.

### Preparation of whole cell lysates

Two approaches of whole cell lysates preparation were used in this work, depending on the scale of the purification.

For large scale purifications, 3 × 10^8^ cells of stable Halo-WDR76 Flp-In™-293, cells were resuspended in 5 x lysis buffer (20 mM HEPES pH 7.5, 0.2% Triton X-100, 1.5 mM MgCl_2_, 0.42M NaCl, 10 mM KCl, 0.5 mM DTT) supplemented with protease inhibitor cocktail (Promega, G6521). Cells were lysed mechanically using a Dounce homogenizer to until more than 90% of nuclei stained positive for trypan blue. Salt active nuclease (ArcticZymes AS, 70900-202) was added to reduce nucleotide-mediated interactions. Whole cell lysates were centrifuged at 400000xg for 30miutes at 4°C to pellet out cell debris from the whole cell lysate (supernatant).

For small scale purifications, 1 × 10^7^ recombinant Flp-In™-293 or HEK293T cells of interest were resuspended in mammalian lysis buffer (50mM Tris-HCl (pH 7.5), 150mM NaCl, 1% Triton® X-100 0.1% Na deoxycholate) supplemented with protease inhibitor cocktail (Promega, G6521). Cells were lysed by mechanically means by passaging of cells through 26-gauge needle 5 times. The resulting lysate was centrifuged at 18,000 × g for 15 minutes at 4 °C and the resulting supernatant collect for downstream applications.

### Native Affinity Purification of WDR76 with associated proteins

For large scale purifications, whole cell lysate or nuclear extracts were diluted in two volumes of lysis buffer (20 mM HEPES pH 7.5, 0.2% Triton X-100, 1.5 mM MgCl_2_, 0.42M NaCl, 10 mM KCl, 0.5 mM DTT) and incubated with pre-equilibrated Magne® HaloTag® beads (Promega) or SNAP-Capture Magnetic Beads (New England Biolabs).

For small scale purifications, lysates were diluted in 700µl of TBS and incubated at 4°C with pre-equilibrated Magne® HaloTag® beads (Promega) or SNAP-Capture Magnetic Beads (New England Biolabs). Bead-lysate mixtures were mixed for overnight at 4°C and washed four times with wash buffer (25 mM Tris·HCl pH 7.4, 137 mM NaCl, 2.7 mM KCl and 0.05% Nonidet® P40). Elution of bound proteins was carried out in 100µl of TEV elution buffer (50 mM Tris·HCl pH 8.0, 0.5 mM EDTA and 0.005 mM DTT containing 2 Units AcTEV™ Protease (Thermo Fisher Scientific/Invitrogen) for 2 hours at 25 °C. In salt resistance assessment, the wash buffer was replaced by high salt buffer (10mM HEPES; pH 7.5, 0.2% Triton x-100, 10mMKCl and 1.5mM MgCl_2_) containing 0.5M, 0.75M or 1.0M NaCl, respectively.

### Gel Filtration chromatography

HaloTag® purified WDR76 protein complexes were prepared from nuclear extracts of stably Halo-WDR76 expressing Flp-In™-293 cells as described above. HaloTag purified WDR76 was loaded onto a Superose 6, 10/300 GL column (Amersham Bioscience) containing the gel filtration buffer (40mM HEPES pH 7.9, 350mM NaCl, 5% (v/v) glycerol, 0.1% (v/v) Tween 20, 1.0mM DTT). The co-fractionation profiles of WDR76 and its binding partners were analyzed by collecting 500µl fractions for identification by LC-MS/MS (MudPIT). The Superose 6, 10/300 GL column was calibrated with the gel filtration markers kit (Sigma-Aldrich cat. # MWGF1000, Blue Dextran 2000 (2000kDa), thyroglobulin (669kDa) Ferritin (440kDa), β-amylase (200kDa), Alcohol dehydrogenase (150), and bovine serum albumin (66kDa).

### Sample preparation for LC-MS analysis

Affinity purified protein isolates were precipitated overnight in 20% trichloroacetic acid at 4°C. Protein precipitates concentrated by centrifugation, washed twice with ice-cold acetone and resuspended in 8M urea tris (pH 8.5) buffer, and 5mM of Tris(2-carboxyethyl) phosphine (TCEP) was added to each sample followed by incubation for 30 minutes followed by addition of 10 of carboxyamidomethylcysteine (CAM) and 30-minute incubation in the dark. Samples were digested in EndoLys-C overnight, followed by additional digestion with Trypsin in 2M urea Tris buffer. Digested samples were loaded onto a 15cm long column with 100µm i.d. × 365μm o.d. fused silica (Polymicro Technologies) packed with 10cm of C18 resin, 4cm strong cation resin and 2cm of C18 resin.

### LC-MS/MS analysis by MudPIT

LC-MS/MS analysis (MudPIT) analysis was performed on Thermo LTQ linear ion trap mass spectrometer (Thermo Fisher Scientific) and peptide separation was carried out using MudPIT. Following MudPIT, RAW files were converted into ms2 files using the RAWDistiller v. 1.0. MS2 files were subjected to database search using ProLuCID algorithm version 1.3.557 and tandem mass compared against human proteins obtained from the National Center for Biotechnology (2016 06 10 release). The database also included common contaminant proteins, including human keratins, IgGs and peoteolytic enzymes and randomized versions of non-redundant protein entry for the estimation of false discovery rate (FDR). NSAF v7 (in-house software) and Q-spec analysis were used to rank proteins identified in each replicate and provide a final report of non-redundant proteins and attribute frequency of appearance (Experiment/control), FDR, log fold change, Z-score to each protein detected in the experiment with respect to background.

### Transfection for confocal Microscopy and Imaging

Flp-In™-293 cells were set at 50% confluency in Makek glass bottom dishes for 24 hours and incubated at 37 °C in 5% CO_2_ for 24 hours. Cells were transfected with 1.5µg of plasmid (expressing Halo-WDR76 and WDR76-Halo) using lipofectamine 2000 according to manufacturer’s instructions. 24 hours post transfection, cells were treated with 20nM TMRDirectTM (stains recombinant Halo-tagged proteins) and cultured at 37 °C in 5% CO_2_ for 24 hours. Cell nuclei were stained with Hoechst for 1 hour at 37 °C in 5% CO_2_. Stained cells were washed twice with warm Opti-MEM® reduced serum medium and imaged with an LSM-700 Falcon confocal microscope.

### Statistical Analysis

Enrichment analysis was performed using QSPEC analysis (Choi et al, 2008) by comparing our purifications relative to mock purifications. We used Z-score≥2, log2FC ≥ 2 or FDR ≤ 0.05 to select the specific proteins in our datasets (Supplementary Tables S1 and S2).

Topological score (i.e. TopS) was applied to the eluted proteins across 26 fractions in the fractionation dataset (Supplementary table S3). We used a TopS cutoff of 5 to select proteins enriched in the 26 fractions. Only proteins identified as well as in the WDR76 wild-type network with TopS greater than 5 in at least one elution were included in the Fig. 6. TopS is written using Shiny application (R package version 3.4.2) for R statistics software. TopS uses several packages, including gplots, devtools and gridExtra. TopS is freely available at https://github.com/WashburnLab/Topological-score-TopS-.

Two TDA networks were constructed using Ayasdi platform as previously used (Lum et al, 2013; Sardiu et al, 2019) Extracting insights from the shape of complex data using topology (Lum et al, 2013). Nodes in the network represent clusters of proteins. We connect two nodes if the corresponding clusters contain a data point in common. The input data for TDA are represented in a bait–prey matrix, with each column corresponding to purification of a bait protein and each row corresponding to a prey protein: values are spectral counts (Fig. 8E) or Z-statistic values (Fig.9E) for each protein. Two types of parameters are needed to generate a topological analysis: First is a measurement of similarity, called metric, which measures the distance between two points in space (i.e. between rows in the data). Second are lenses, which are real valued functions on the data points. Variance normalized euclidean with Neighborhood lens 1 and Neighborhood lens 2 (resolution 30, gain 3.0x) were used to generate Fig. 8E and variance normalized euclidean metric with PCA1 and PCA2 (resolution 30, gain 3.0x) was used to generate Fig 9E.

## Data Availability

Affinity purification coupled mass spectrometry data is available via massive MassIVE ID #MSV000083778 (username: Dayebga_1 with the password: baligashu) via the link https://massive.ucsd.edu/ProteoSAFe/dataset.jsp?task=c2442c560afa4c3980d064eb263e2621

## Acknowledgements

Research reported in this publication was supported by the Stowers Institute for Medical Research and the National Institute of General Medical Sciences of the National Institutes of Health under Award Number RO1GM112639 to MPW. The content is solely the responsibility of the authors and does not necessarily represent the official views of the National Institutes of Health.

## Author Contributions

G.D., M.E.S., L.F., and M.P.W. designed the experiments. G.D. performed experiments. G.D. and M.E.S. performed computational analyses of data. G.D., M.E.S, L.F., and M.P.W. wrote the manuscript.

