## Supplemental Information and Supplemental Figures for "Biochemical Reduction of the Topology of the Diverse WDR76 Protein Interactome"

**Index of Supplementary Data**

**Primer sequences:**

1.Primers for construction of WDR76-Halo tagged pcDNA5/FRT expression plasmid

Forward primer (PacI_WDR76): CTA TAG GGA GAC CCA AGC TGT TAA TTA ACA TGT CCA GGT CGG GCG CG

Reverse primer (NheI_WDR76): TGGTTG GCT CGA GAG AAA CGC TAG CGC AGC TTT TTT CAT TCA TAA AAA CAT GTA TCT T

2. Primers for construction of pcDNA5/FRT construct for expression of SNAP-tagged WDR76Δ (aa 1-310) deletion mutant.

Forward primer (WDR76_trunc_310_F): GTC ATT AGT GAA GAT ACC GTT TAC AAA TAA GTT ACC ACA GGC CCA ATA TTC TCT ATG

Reverse primer (WDR76_trunc_310_R): GCC ATA GAG AAT ATT GGG CCT GTG GTA ACT TAT TTG TAA ACG GTA TCT TCA CTA ATG A

3. Primers for construction of pcDNA5/FRT construct for expression of SNAP-tagged WDR76Δ’ (aa 311-626).

Forward primer (WDR76_trunc_WD40F): GAC GAT GAT GAC AAG GCG ATC GTT ACC ACA GGC CCA ATA TTC

Reverse primer (WDR76_trunc_WD40R):

CGA GGC TGA TCA GCG GGT TTT CAG CAG CTT TTT TCA TTC ATA AAA AC

**Supplementary Figure Legends**

**Figure S1. Schematic of 17 unique WDR76 homologous sequences available in Homologene (NCBI HomoloGene:38573).** We represented a cartoon of all the proteins to show the presence of the conserved C-terminal WD40 repeat domain (colored box).

**Figure S2. Localization of Halo-WDR76 and WDR76-Halo in HEK293FRT cells.** AP-MS analysis showed depth of the WDR76 interaction with Halo-WDR76 (N-terminal HaloTag) compared to WDR76-Halo (C-terminal HaloTag). (A) Schematic representation of Halo-WDR76 and WDR76-Halo. Note: Following TEV cleavage, WDR76 is eluted leaving behind the HaloTag. (B) Whole cell lysate of 293FRT cells transiently transfected with empty Halo vector (lane 2), Halo-WDR76 vector (lane 3) and WDR76-Halo vector (lane 4) were prepared as detailed in materials and methods. The lysates were subjected to SDS-PAGE and western blot analysis using the anti-HaloTag® polyclonal antibody (with anti-tubulin antibody as a loading control). (C-D) Localization of Halo-WDR76 and WDR76-Halo in 293FRT cells, respectively. Cell nuclei were stained with Hoechst (blue) and recombinant Halo-WDR76 or WDR76-Halo with TMRDirect ligand (red). (E-F) Wider view and three panels of localization of Halo-WDR76 and WDR76-Halo in 293FRT cells, respectively. Cell nuclei were stained with Hoechst (blue) and recombinant Halo-WDR76 or WDR76-Halo with TMRDirect ligand (red).

**Figure S3. High confidence interactome of WDR76 at 0.42M NaCl purification buffer.** (A) Scatter plot of AP-MS results from chromatin-enriched nuclear extracts of Halo-WDR76 stable cells. (B) Scatter plot of AP-MS results from whole cell extracts prepared from Halo-WDR76 stable cells. (C) Schematic of the workflow used in Gilmore et al.

**Figure S4. Functional annotations of salt-resistant WDR76 interactions in stable Halo-WDR76 expressing HEK293FRT cells.** (A) CORUM analysis of WDR76 which persist at

NaCl between 0.5 and 1.0. (B) Gene ontology (GO) analysis of the biological processes enriched at 1.0 salt concentrations.

**Description of Supplementary Tables**

**Table S1. AP-MS analysis of WDR76 interactomes in Halo-WDR76 and WDR76-Halo expresses in HEK293FRT cells.** Here we present the Q-spec results for Halo-WDR76 (A) and WDR76-Halo (B). We also present protein list of the proteins with z-score greater than 2 and FDR value of less than 0.05 in the Halo-WDR76 purification (C) and WDR76-Halo purification (D).

**Table S2**. **High-confidence AP-MS datasets from 293FRT cells with stable expression of Halo-WDR76 (0.42M NaCl purification buffer)**. Here we present the Q-spec data for WDR76 AP-MS analysis published in Gilmore et al (A) and protein list of proteins with z-score greater than 2 and (log fold change greater than 2 or FDR less than 0.05) (B). (C) Q-spec analysis of AP-MS data from chromatin-enriched nuclear extract. (D) List of proteins with z-score greater than 2 and log fold change greater than 2 or FDR less than 0.05 in chromatin-enriched nuclear extract data. (E) Q-spec analysis of AP-MS data from whole cell extract. (F) List of proteins with z-score greater than 2 and log fold change greater than 2 or FDR less than 0.05 in whole cell extract AP-MS data. (E) Original output of cellular components from downloaded from gprofiler. (F) List of protein lists in each category GO category.

**Table S3. Size exclusion analysis of WDR76 isolates on a superose 6 column** (A) Proteins identified by affinity purification coupled with size exclusion coupled mass spectrometry of WDR76 isolates. (B) Topological score (i.e. TopS) of the proteins detected across the 26 fractions analyzed. (C) A representative subset of (B) showing the topological scores of a subset of important proteins identified. Next, we present Gene Ontology (GO) terms: Biological processes enriched in both low and high molecular weight WDR76 interactions (D) only low molecular weight interactions (E) and high molecular weight interactions (F).

**Table S4. Domain-specific interactions of WDR76. (A)** Q-spec results between WDR76 purifications for full length WDR76 and WDR76 deletion: WDR76Δ (1-310) and WDR76Δ' (311-626). (B) Q-spec results between WDR76 purifications for full length WDR76 and WDR76 deletion: WDR76 (1-310) and WDR76 (311-626). Subset of proteins that change in at least one deletion.

**Table S5. Evaluation of the salt resistance of the WDR76 interactome.** (A) Q-spec analysis of AP-MS data for WDR76 at 0.5, 0.75 and 1.0 NaCl wash conditions. (B) Average Proteins abundance represented by dNSAF for proteins with Z-score greater than 2 and log fold change of greater than 2 or FDR less than 0.05 for the for 0.5, 0.75 and 1.0 NaCl wash conditions**.** (C) Topological scores were calculated for proteins passing a statistical criterion in worksheet 2. (D) Q-spec analysis of WDR76 interactions at 1.0M NaCl wash conditions, respectively (Z-score ≥2, (log fold change≥2 or FDR≤0.05)). (E) CORUM analysis of proteins passing the criteria in 6D.

**Table S6. Reciprocal AP-MS validation of WDR76 interaction GAN, HELLS and SIRT1 by transient transfections in 293FRT cells** (A) Q-spec analysis of AP-MS data for Halo-WDR76, Halo-GAN, Halo-HELLS and Halo-SIRT1 in HEK 293FRT cells. (B) Subset of proteins from A with z-score greater than 2and FDR less than 0.05. (C) Q-spec results for reciprocal AP-MS for WDR76, GAN, HELLS and SIRT1. Here are proteins that have a Z-score greater than 2 and an FDR less than 0.05.

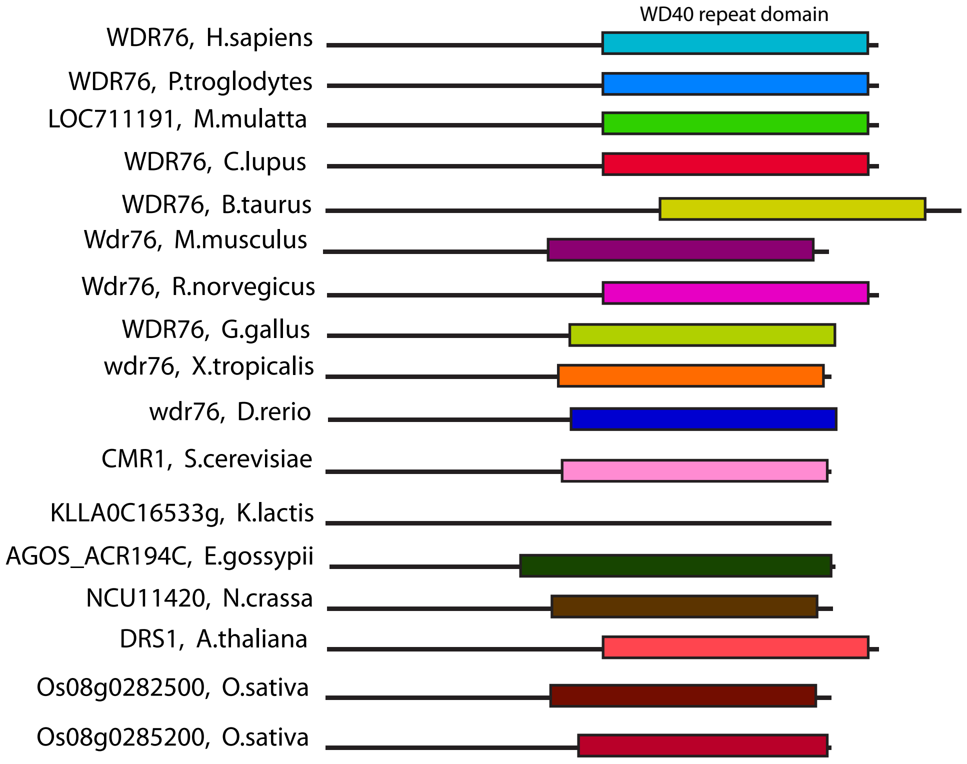


Supplementary Figure 1. Dayebgadoh et al.

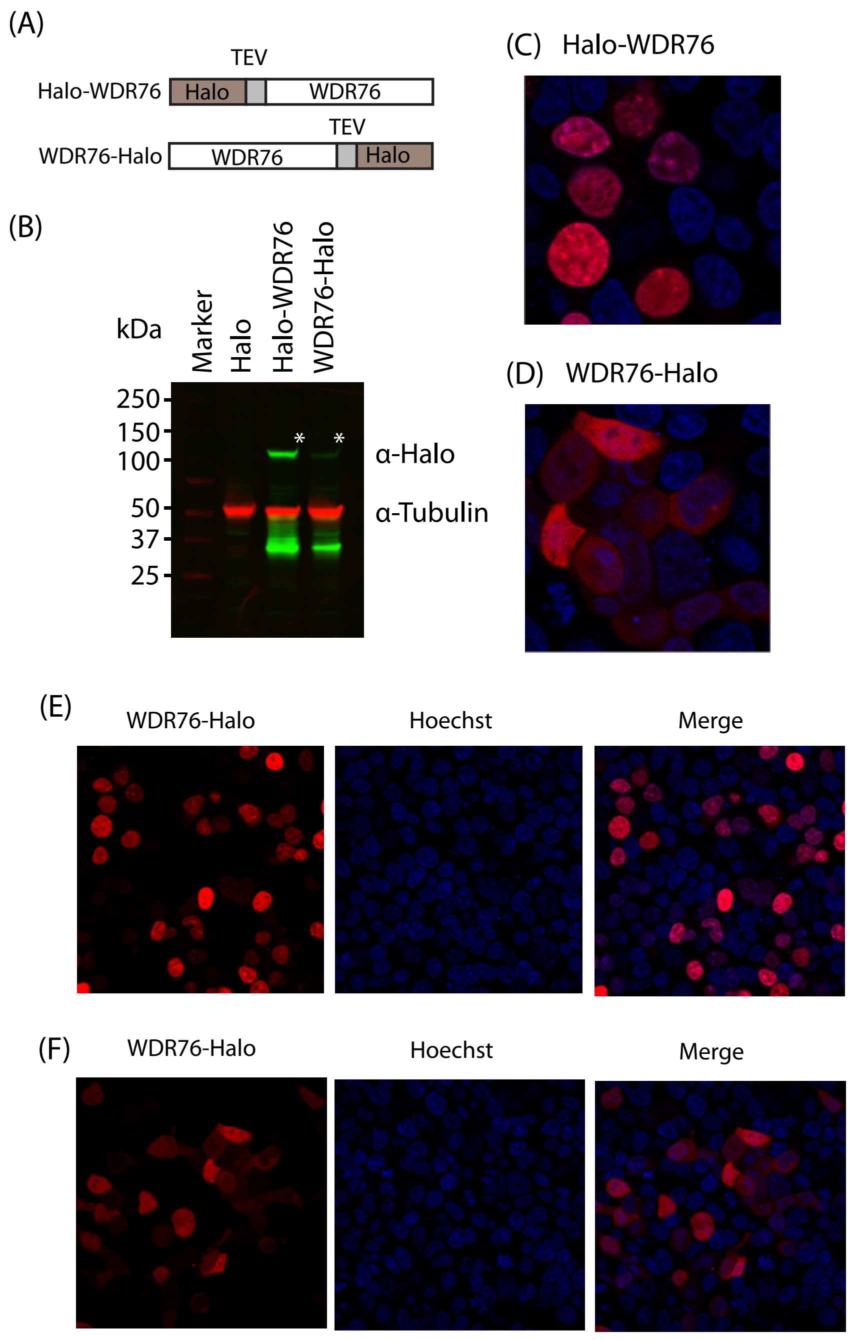


Supplementary Figure 2. Dayebgadoh et al.

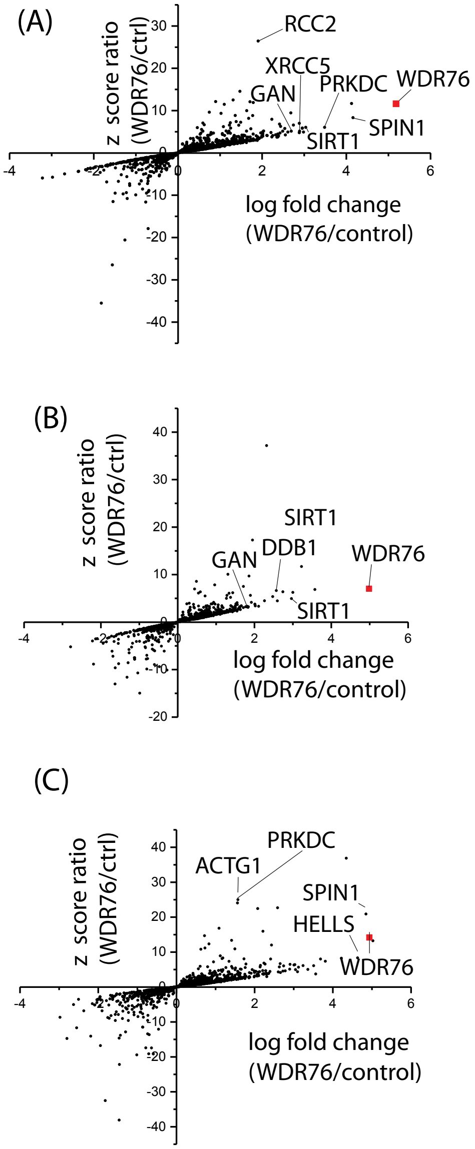


Supplementary Figure 3. Dayebgadoh et al.

(A)

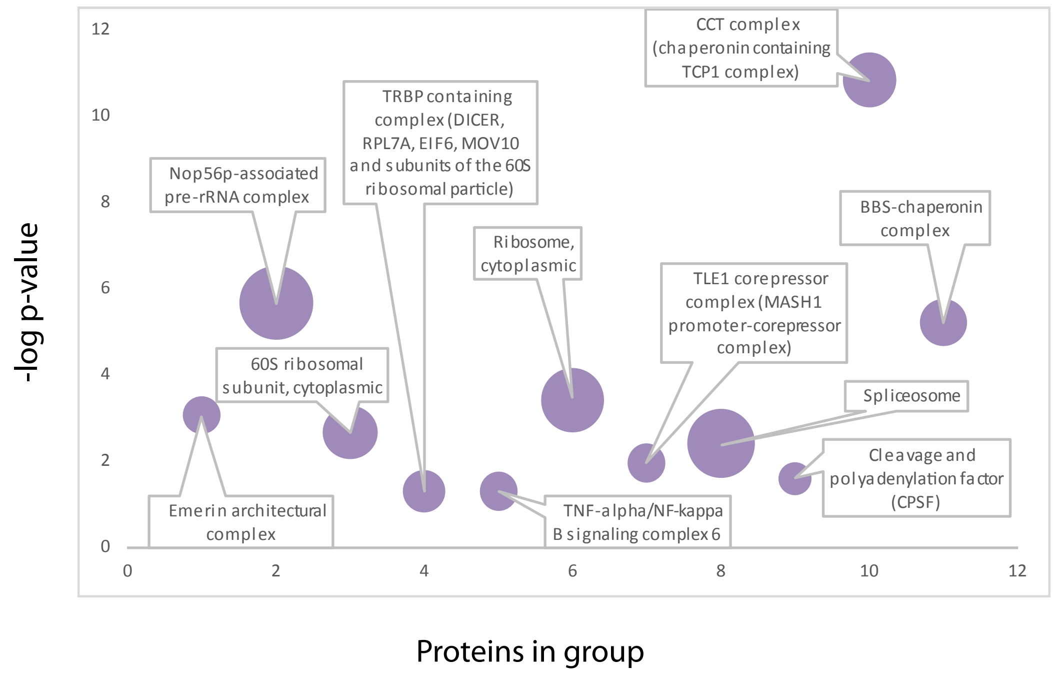


(B)

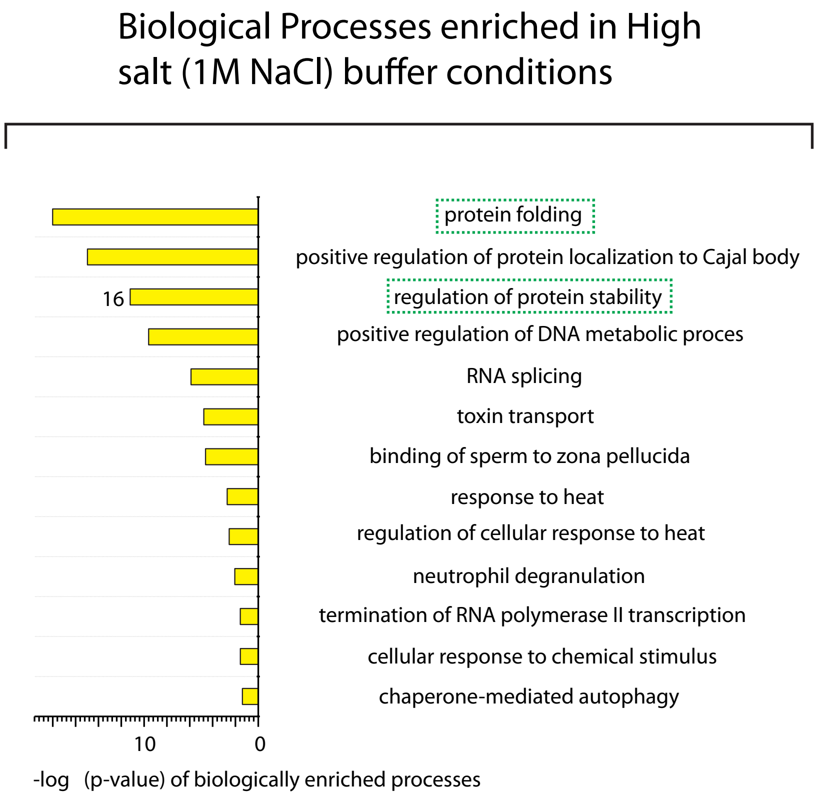


Supplementary Figure 4. Dayebgadoh et al.
